# PITX2C Deficiency Promotes Arrhythmogenic Remodeling via Oxidative Stress in Atrial Myocytes

**DOI:** 10.64898/2026.03.27.714813

**Authors:** Andy Kim, Sébastien Gauvrit, Frederick S. Vizeacoumar, Michelle M. Collins

**Affiliations:** Department of Anatomy, Physiology, and Pharmacology, College of Medicine, University of Saskatchewan, 107 Wiggins Road, Saskatoon, SK, S7N 5E5; Department of Pathology and Laboratory Medicine, College of Medicine, University of Saskatchewan, 107 Wiggins Road, Saskatoon, SK, S7N 5E5

**Keywords:** Atrial fibrillation, PITX2, Neonatal rat atrial myocyte, Mitochondrial dysfunction, Reactive oxygen species

## Abstract

**Aims:** Genome-wide association studies have identified numerous cardiac transcription factors in association with atrial fibrillation. Amongst these transcription factors, the paired-like homeodomain transcription factor 2 (*PITX2*) is the strongest genetic risk variant associated with atrial fibrillation. However, the downstream mechanisms of PITX2 are not completely understood. Here, we explore the role of PITX2 in oxidative metabolism and stress as a unifying mechanism of arrhythmogenesis.

**Methods and results:** To identify PITX2 mechanisms, we performed transcriptomic analysis in *Pitx2c*-deficient neonatal rat atrial myocytes. We identify oxidative phosphorylation as the top dysregulated pathway and direct transcriptional targets lie in mitochondrial electron transport chain complexes I and IV. Using the Seahorse Extracellular Flux Analyzer, we identified a functional decrease in oxidative metabolism in *Pitx2c*-deficient cardiomyocytes. As electron transport chain complexes I and IV may generate reactive oxygen species (ROS) under mitochondrial dysfunction, we quantified mitochondrial specific ROS using MitoSOX and observed an increase in mitochondrial specific ROS in *Pitx2c*-deficient cardiomyocytes. We additionally assessed spontaneous cardiomyocyte calcium cycling using Fluo-8AM and observed an increased frequency of pro-arrhythmogenic mechanisms including early and delayed afterdepolarizations as inferred through calcium traces. Further, we identified sarcomere disassembly including a potential role of PITX2 in regulating Titin, where *Pitx2c*-deficient cardiomyocytes display Titin mis-localization within the sarcomeres. To assess whether ROS drives these phenotypes, we treated neonatal rat atrial myocytes with N-acetylcysteine, a potent ROS scavenger, and observed decreased early and delayed afterdepolarizations, as well as restoration of Titin localization.

**Conclusion:** PITX2C maintains atrial metabolism and redox balance; the loss of PITX2C results in reduced oxidative metabolism and an elevation in oxidative stress that ramifies cardiomyocyte dysfunction. Treatment with antioxidant restores AF-associated phenotypes including abnormal calcium cycling and sarcomere disassembly in *Pitx2c*-deficient atrial cardiomyocytes.

**TRANSLATIONAL PERSPECTIVE:** Genetic variants close to the *PITX2* gene associate most strongly with atrial fibrillation. This study reveals a mechanistic link between multiple AF-associated phenotypes and mitochondrial dysfunction with subsequent accumulation of reactive oxygen species downstream of PITX2. Importantly, metabolic therapies and reducing oxidative stress may present a potential clinical strategy to reverse and prevent functional and structural remodelling related to AF.

## 1. INTRODUCTION

Atrial fibrillation (AF) is the most common cardiac arrhythmia. Genome wide association studies have identified genetic variants on chromosome 4q25, near the paired-like homeodomain transcription factor 2 (PITX2) gene, as the most significantly associated with AF(Roselli et al., 2018; Van Ouwerkerk et al., 2019, 2020). Animal models of *PITX2* insufficiency have highlighted its critical role in AF pathogenesis. Studies in *Pitx2* heterozygous mice have shown that PITX2 has a transcriptional role in maintaining cardiac electrical stability and structure with targets in ion channels, calcium handling, and intercalated disk formation(Y. Tao et al., 2014; Tarifa et al., 2023). Similarly, we have previously demonstrated that *pitx2c* deficient zebrafish develop sarcomere and metabolic defects that alter cardiac function in early larval stages and which contribute to electrical instability and structural remodelling in adult zebrafish(Collins et al., 2019). Although there is a clear role of PITX2 in maintaining components of cardiac rhythm, how these components are linked is incompletely understood.

One central mechanism of PITX2 in AF pathogenesis we propose is through its transcriptional regulation of oxidative metabolism and stress(G. Tao et al., 2016). It has been previously demonstrated that *pitx2c*-deficient zebrafish have reduced oxidative phosphorylation capacity and Pitx2 transcriptionally regulates redox genes(Collins et al., 2019). Additionally, human induced pluripotent stem cell (hiPSC) *PITX2* knockout models have shown that *PITX2* deficiency alters atrial metabolism resulting in a metabolic shift towards glycolysis that contributes to structural and electrical remodelling(Reyat et al., 2024). Recently, reactive oxygen species (ROS) have been demonstrated as a unifying mechanism of AF susceptibility in *Pitx2* insufficient mice; pharmacological targeting of ROS-generated lipid dicarbonyls prevented arrhythmia vulnerability in *Pitx2* insufficient mice(Subati et al., 2025). However, an interesting question that remains is whether PITX2-dependent alterations in intracellular calcium dynamics are driven by oxidative stress. Therefore, to further support this unifying mechanism, we characterized the role of ROS in *Pitx2* deficiency in neonatal rat atrial myocytes. We demonstrate that reduction of ROS ablates arrhythmia susceptibility, restoring aberrant calcium cycling and structural abnormalities.

## 2 MATERIALS AND METHODS

### 2.1 Isolation of neonatal rat atrial myocytes (NRAM)

All protocols were in accordance with the guidelines detailed by the Canadian Council of Animal Care by the Animal Research Ethics Board at the University of Saskatchewan (AUP #20220090).

This protocol was adapted from Wu et al(Wu et al., 2020). 2-day old Sprague Dawley (Charles River) neonatal rat pups were euthanized by decapitation and the heart was harvested in PBS supplemented with 5mM glucose. Ventricles and large vessels were removed, and atria were subjected to enzymatic digestion using a lysis buffer solution containing 20mM HEPES-NaOH, 130mM NaCl, 3mM KCl, 1mM NaH_2_PO_4_, 5mM glucose, 15mg/mL DNase I (Roche 104159), collagenase Type II (ThermoFisher 17101015), and pancreatin (Sigma P3292). Multiple rounds of enzymatic digestion were performed where the tissue was incubated with 10mL lysis buffer at 37°C for 10 minutes. Supernatant from the first round of digestion was discarded and subsequent rounds were performed with the supernatant harvested in 5mL of horse serum. Once the tissue was completely digested, cells were pelleted at 300 x *g* for 5 minutes at 4°C. The cell pellet was then re-suspended in pre-warmed red blood cell lysis buffer (Sigma 11814389001) and incubated for 3 minutes at 37°C after which, 20mL of DMEM/F-12 (ThermoFisher 11320033) supplemented with 10% FBS and 1% Penicillin-Streptomycin. Cells were pelleted at 300 x *g* for 5 minutes at 4°C. The cell pellet was then again re-suspended in DMEM/F-12 with 10% FBS and 1% Penicillin-Streptomycin and plated on uncoated cell culture dishes for 90 minutes to remove fibroblasts and endothelial cells. Unadhered cells were then collected, passed through a 70μm mesh filter, centrifuged at 300 x *g* for 5 minutes at 4°C, re-suspended in culture medium (DMEM/F-12, 10% FBS, 1% Penicillin-Streptomycin, 5% horse serum, 3mM Na-pyruvate, 0.1mM ascorbic acid, 1:200 insulin/transferrin/Na-Selenite solution, 0.2% BSA, and 20uM cytarabine (Sigma C1768). Cells were seeded at a cell density of 60 × 10^4^ cells/cm^2^. The medium was replaced the next day to remove dead cells. To assess the effects of oxidative stress, cells were treated using N-acetylcysteine (NAC) at 1mM for 24 hours prior to experiments.

### 2.2 *Pitx2c* siRNA transfection

Cells were left to reach 60-70% confluency and transfected with 25nM, 50nM, and 100nM non-targeting siRNA (Invitrogen 4390843), *Pitx2c* s132548 siRNA (Invitrogen 439077) or *Pitx2c* s132549 siRNA (Invitrogen 439077) diluted in OptiMEM (ThermoFisher 31985070) using Lipofectamine RNAiMAX (ThermoFisher 13778150) in accordance with the manufacturer’s protocol. After 24 hours, the transfection medium changed into the culture medium stated above. Cells were harvested at 24, 48, and 72 hours post transfection to determine the time of maximal *Pitx2c* knockdown. Cells were then pelleted and stored at -80°C until RNA extraction was performed.

### 2.3 RNA Extraction and reverse-transcriptase-qPCR

RNA extraction was performed using the Qiagen RNeasy Plus Micro Kit (Qiagen 74034) in accordance with the manufacturer’s protocol. RNA concentration and A_260_/_A280_ ratios was assessed by spectrophotometry (Nanodrop One). First strand cDNA was synthesized using Maxima First Strand cDNA Synthesis Kit for RT-qPCR with dsDNase (ThermoFisher K1641). Gene knockdown was confirmed by RT-qPCR using *ACTB* (B-actin) as the reference gene. Each qPCR reaction was 10µL containing 1X PowerUp™ SYBR™ Green Master Mix (ThermoFisher A25742), 0.2µM forward primer, 0.2µM reverse primer and 2µL template cDNA. To analyze the fold change of the genes of interest, the delta-delta Ct method was applied which uses the formula 2^-ΔΔCt^ where ΔΔCt = ΔCt (treated sample) – ΔCt (untreated sample) and ΔCt = Ct (gene of interest) -Ct (reference gene). Primer sequences used for RT-qPCR are provided in Supplemental Table 1.

### 2.4 RNA-sequencing

Total RNA was isolated from quadruplicates of *Pitx2c*-deficient neonatal rat atrial myocytes and their controls. RNA yield and quality were assessed and samples with sufficient RNA quality (RNA integrity number > 8) were sequenced. Poly A-selected libraries were sequenced by the Centre for Applied Genomics at the University of Toronto and mapped to the rat GRCr8 genome. Quality control was performed using FastQC v.0.12.1. Differential gene expression analysis was performed with DESeq2 v.1.26.0 and principal component analysis performed. Significant differentially expressed genes were classified as log2FoldChange> +/-0.85 (1.5 fold change) and p-adj <0.05. Gene set enrichment analysis was then performed to identify global changes in gene expression patterns using the Hallmark gene set. Significantly enriched pathways were classified as false discovery rate <0.25.

### 2.5 Identification of genes associated with ChIP-seq peaks

ChIP-seq peaks identified in GSM116257(Y. Tao et al., 2014) were obtained from the BED file. Peaks were annotated to the nearest gene using GENCODE mouse genome annotation (GRCm39/mm39). Gene annotations were obtained in GTF format and converted to transcript database objects using the GenomicFeatures package in R. Peak-to-gene associations were performed using ChIPseeker (R library), assigning each peak to the closest transcription start site (TSS) within a ±3 kb window; peaks located outside this window were annotated to the nearest gene based on genomic distance. Gene identifiers were mapped to official mouse gene symbols using org.Mm.eg.db. A non-redundant list of genes associated with ChIP-seq peaks was generated for downstream analyses.

### 2.6 Seahorse XFp

Seahorse mitochondrial stress test (Agilent 103010-100) and the glycolytic rate assay (Agilent 103346-100) were performed on neonatal rat atrial myocytes seeded into 8-well XFp microplates at a density of 40 000 cells/well.

#### 2.6.1 MitoStress test

Cells were washed with XF DMEM medium (Agilent 103575-100) supplemented with 1mM pyruvate, 2mM glutamine, and 10mM glucose and placed in a 37°C without CO_2_ for 45 minutes. An automated Seahorse XFp protocol was performed consisting of an initial calibration/equilibration, followed by injection of drugs. For the mitochondrial stress test, ports were loaded with (A) 2μM oligomycin, (B) 2μM FCCP, and (C) 0.5μM rotenone/antimycin A.

#### 2.6.2 Glycolytic rate assay

Cells were washed with XF DMEM medium (Agilent 103575-100) supplemented with 1mM pyruvate, 2mM glutamine, and 10mM glucose and placed in a 37°C without CO_2_ for 45 minutes. An automated Seahorse XFp protocol was performed consisting of an initial calibration/equilibration, followed by injection of drugs. For the glycolytic rate assay, ports were loaded with (A) 0.5μM rotenone/antimycin A and (B) 2-deoxy-D-glucose (2-DG).

### 2.7 Mitochondrial DNA content

Genomic DNA (gDNA) was isolated extracted using the Qiagen DNeasy Blood and Tissue Kit (Qiagen 69506). Relative mitochondrial DNA (mDNA) copy number was measured by a qPCR-based method to calculate the ratio of mtDNA/nDNA. mtDNA was quantified using primers for *mt-co1* and nDNA was quantified using primers for *polg*. Each qPCR reaction was 10µL containing 1X PowerUp™ SYBR™ Green Master Mix, 0.2µM forward primer, 0.2µM reverse primer and 3µL gDNA. To analyze the fold change, the delta-delta Ct method was applied which uses the formula 2^-ΔΔCt^ where ΔΔCt = ΔCt (treated sample) – ΔCt (untreated sample) and ΔCt = Ct (*mt-co1*) -Ct (*polg*). Primer sequences used for qPCR are provided in Supplemental Table 1.

### 2.8 MitoSOX

Neonatal rat atrial myocytes were loaded with 0.5μM MitoSOX Red (ThermoFisher M3600) diluted in Opti-MEM. The sample was incubated at 37°C for 30 minutes after which the cells were washed 3 times with pre-warmed Opti-MEM. Imaging was performed with the Nikon Eclipse Ti2 CSU-X1 using a 60x objective and cells were maintained at 37°C with 5% CO_2_ while imaging. Images were analyzed from three regions of interest (50 px x 50px) within the cytosol and taking the mean of these measurements for each cardiomyocyte using Fiji(Schindelin et al., 2012) software.

### 2.9 Fluo-8AM calcium imaging

Neonatal rat atrial myocytes were loaded with 5µM of the calcium indicator Fluo-8 AM (ab142773) diluted in Opti-MEM. The sample was incubated for 90 minutes at 37°C. After loading, the cells were washed with Opti-MEM for 5 minutes 3 times. Calcium imaging was performed with the Nikon eclipse Ti2 using a 60x objective. Movies were taken with a size of 1192 × 1192 pixels for 15 seconds with 50 ms delay.

Calcium analysis was performed in ImageJ by drawing a region of interest in the cytosol of the cell and fluorescence values were exported to OriginLabs for analysis using the calcium transient analysis app(Hawey et al., 2023).

### 2.10 Immunocytochemistry

Neonatal rat atrial myocytes cultured in Ibidi dishes (Ibidi 80807) were washed in PBS, fixed in PEM fixative (3% paraformaldehyde (PFA), 100mM Pipes, 1mM MgSO_4_, and 2mM EGTA in distilled water and adjusted to pH 7.4) for 15 minutes at room temperature, then washed with ice cold PBS, permeabilized with 0.1% Triton X-100 in PBS for 30 minutes at room temperature, and blocked with 10% sheep serum in PBS for 1 hour at room temperature. Primary antibody labelling was performed overnight at 4°C, washed with PBS, and labelled with secondary antibody for 1 hour at room temperature. Images were acquired using a Nikon Eclipse Ti2 CSU-X1 using a 40x objective. Primary antibodies used were alpha-Actinin (1:100, Sigma A7732), Myomesin (1:20, DSHB mMac) and Titin (1:100, DHSB 9 D10). Secondary antibody used was anti-mouse 647 (1:500).

Images were analyzed in Fiji software for cellular morphology, PatternJ(Baheux Blin et al., 2024) plugin for Titin co-localization, and Ridge Detection(Steger, 1998; Wagner et al., 2017) plugin for sarcomere density. Sarcomere length and organization was analyzed using SotaTool(Stein et al., 2022).

### 2.11 Statistical Analysis

Statistical analyses were performed using GraphPad Prism 10. The specific statistical tests used are reported in the figure legends. Results were considered to be statistically significant if the probability (*p*) value was <0.05 as the threshold to reject the null hypothesis.

## 3 RESULTS

### 3.1 The PITX2C gene regulatory network in the post-natal atria

To assess the gene regulatory network downstream of PITX2 in the post-natal heart, we leveraged neonatal rat atrial myocytes (NRAMs) transfected with *Pitx2* siRNA. Significant downregulation of *Pitx2c* mRNA (86.03%) was observed at 48 hours post transfection compared to control siRNA treated cells (Figure S1). Poly(A) bulk RNA-seq was then performed to identify PITX2-dependent gene expression changes. Principal component analysis identified close and distinct clustering of both *Pitx2c* deficient and control cardiomyocytes (Figure 1A). 230 transcripts showed significant differential expression, including significant downregulation of *Pitx2* (Figure 1B). Gene set enrichment analysis identified oxidative phosphorylation, fatty acid metabolism, and glycolysis amongst the top dysregulated pathways, suggesting a metabolic role of PITX2 (Figure 1C). Transcripts enriched in oxidative phosphorylation encode mitochondrial electron transport chain proteins including complex I and IV subunits and ATP synthase subunits (Figure 1D). To determine whether these changes are direct transcriptional targets of PITX2, we analyzed a previously published ChIP-seq dataset performed in 12-week-old mouse heart tissue(Y. Tao et al., 2014). We identify direct PITX2 targets that encode complex I and IV subunits and ATP synthase subunits exist within the mouse heart suggesting a direct transcriptional role of PITX2 in regulating metabolic processes and oxidative stress (Table S2).

**Figure 1.**
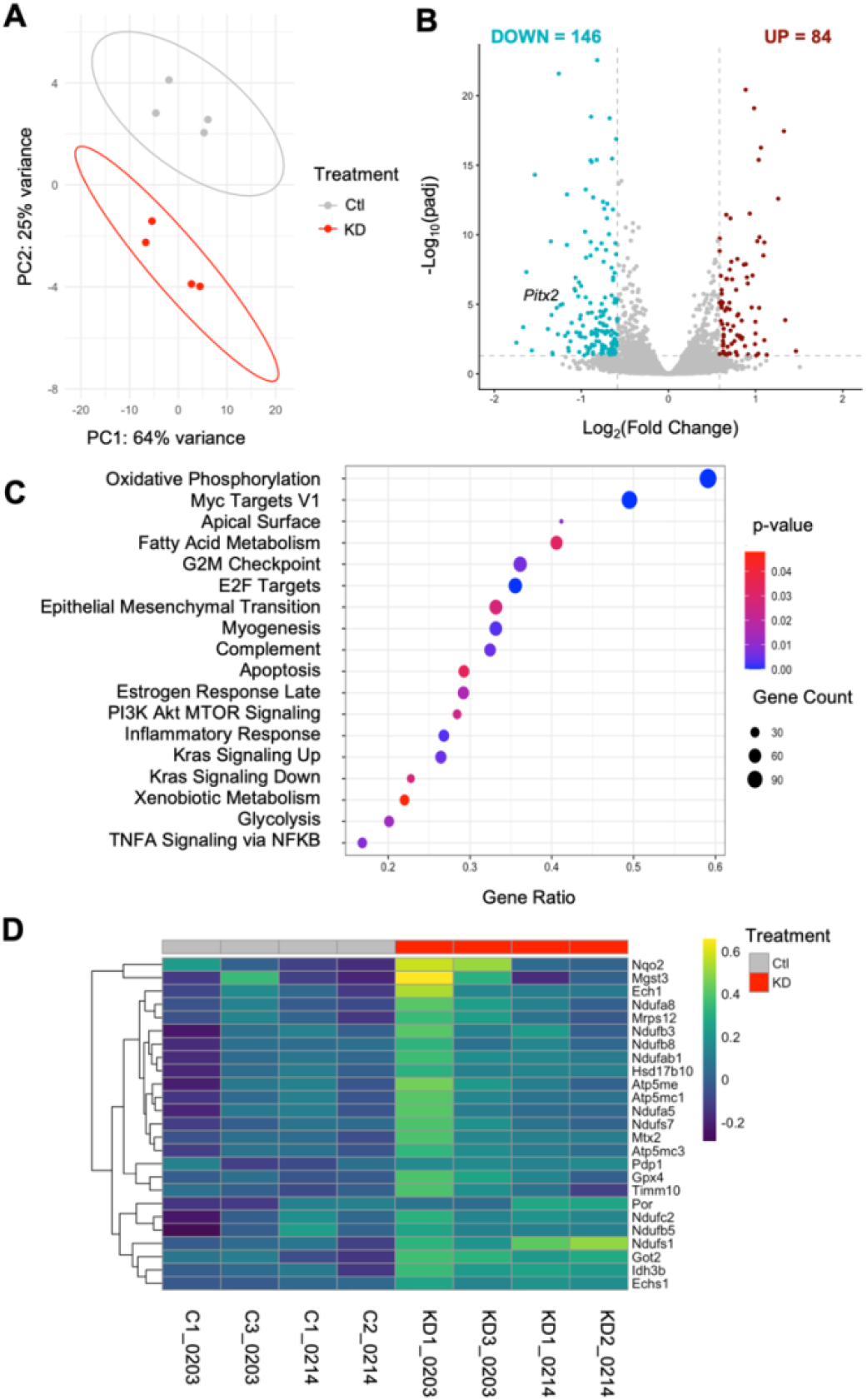
PITX2C transcriptionally regulates oxidative phosphorylation pathway genes. Bulk RNA-seq was performed on neonatal rat atrial myocytes (NRAMs) transfected with non-targeting (Ctl) or *Pitx2c* (KD) siRNA. [A] Principal component analysis of samples used in RNA-seq with ellipses depicting 95% confidence intervals. [B] Volcano plot showing significantly (p<0.05) enriched genes with at least a |1.5| fold change. [C] Gene set enrichment analysis (GSEA) of significantly (FDR >0.25) enriched pathways. [D] Top 25 genes enriched in oxidative phosphorylation pathway from GSEA.

### 3.2 PITX2C regulates atrial cardiomyocyte metabolism and oxidative stress

To evaluate if the PITX2C-induced dysregulation of mitochondrial oxidative metabolism gene expression translated to abnormal mitochondrial bioenergetics, we utilized the XFp Extracellular Flux Analyzer to measure oxygen consumption rate (OCR) as an indirect quantification of oxidative phosphorylation. OCR was assessed at multiple time points before and after the addition of different inhibitors (Figure 2A) of the mitochondrial electron transport chain which allow for the evaluation of OCR based parameters. *Pitx2c*-deficient NRAMs exhibit decreased non-mitochondrial and basal respiration in comparison to control cells (Figure 2B). The addition of oligomycin (an ATP-synthase inhibitor) allows us to distinguish whether the changes in basal OCR are subsequent of decreases in ATP-linked mitochondrial OCR or non-ATP linked OCR as a secondary consequence to electron transport chain proton leak. We observe no difference between control and *Pitx2c*-deficient NRAMs indicating the decrease in basal OCR was primarily due to changes in ATP-linked OCR (Figure 2B). Following, FCCP (electron transport chain uncoupler) was injected to mimic heightened energy demands inducing maximal respiration capacity to reduce oxygen into water, as a component of normal respiration. *Pitx2c*-deficient cardiomyocytes show decreased maximal respiration and spare respiratory capacity (Figure 2B). To assess whether this decrease in mitochondrial respiration was due to changes in cellular mitochondrial content, we quantified the mitochondrial DNA (mtDNA) content and observed no change between control and *Pitx2c*-deficient cardiomyocytes (Figure 2C). In addition to oxidative phosphorylation, glycolysis contributes to energy production within the post-natal heart and metabolic switches to glycolysis is observed when oxidative phosphorylation is reduced to compensate for decreased mitochondrial energy production. Therefore, we assessed glycolytic function through the proton efflux rate (PPR) (Figure S3A). We observe no significant changes in basal glycolysis and compensatory glycolysis with the injection of rotenone and antimycin A (electron transport chain complex I and III inhibitor) in *Pitx2c*-deficient NRAMs (Figure S3B).

**Figure 2.**
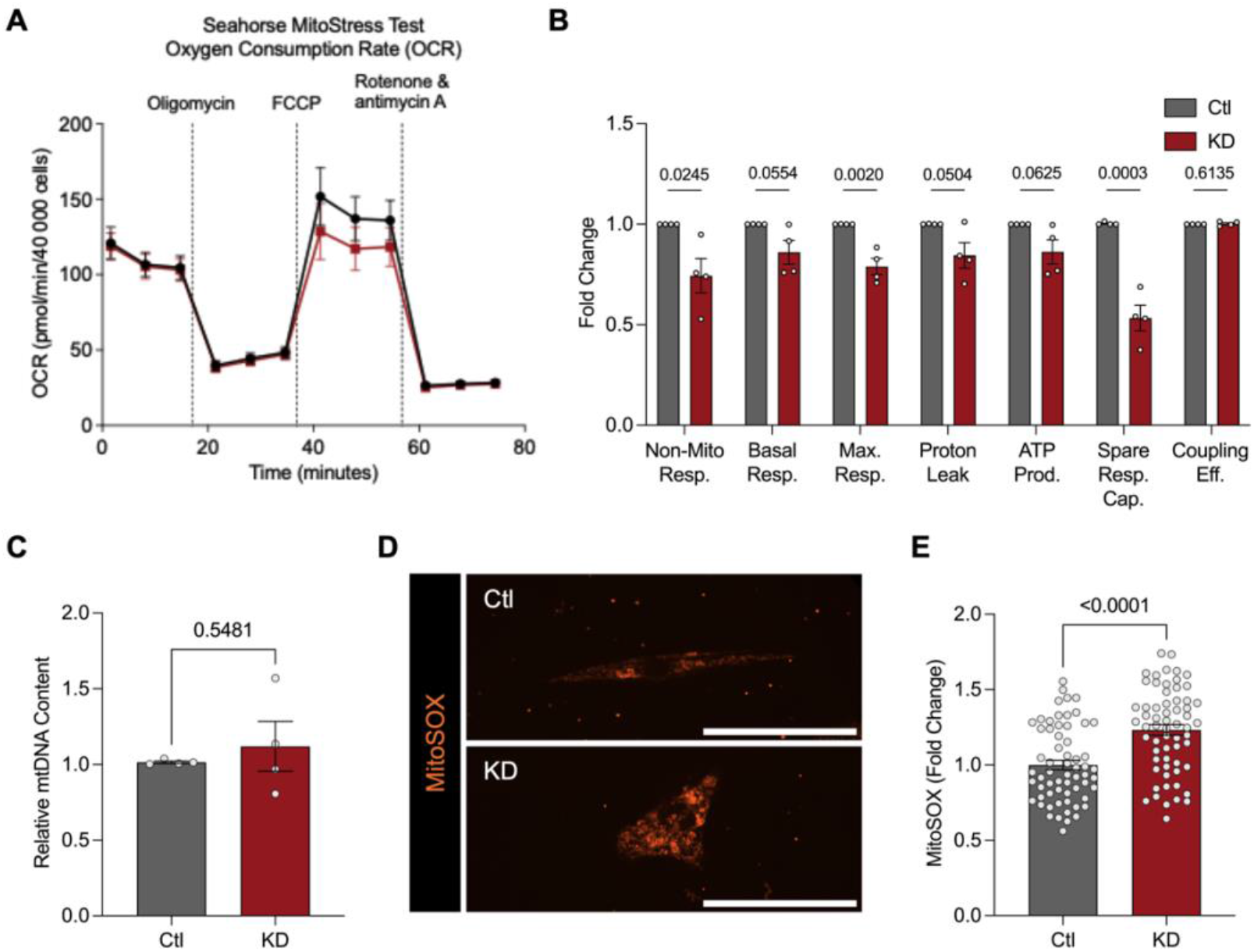
*Pitx2c* deficiency impairs mitochondrial oxidative phosphorylation and elevates mitochondrial derived reactive oxygen species. Mitochondrial respiration was assessed in neonatal rat atrial myocytes (NRAMs) transfected with non-targeting (Ctl) or *Pitx2c* (KD) siRNA using the Seahorse XFp MitoStress test. [A] Representative oxygen consumption rate (OCR) trace is shown with injections of oligomycin, carbonyl cyanite-4 (trifluoromethoxy) phenylhydrazone (FCCP), and rotenone/antimycin A. [B] Quantification of mitochondrial function relative to Ctl cells including non-mitochondrial respiration (Non-Mito Resp.), basal respiration (Basal Resp.), maximal respiration (Max Resp.), Proton Leak, ATP production (ATP Prod.), spare respiratory capacity (Spare Resp. Cap.), and coupling efficiency (Coupling Eff.). n=4. [C] mtDNA content relative to Ctl cells was quantified by qPCR and are expressed as *mt-co1*/*polg* (mtDNA/nDNA). n=4. Data are expressed as means± SEM. [D] Mitochondrial specific reactive oxygen species accumulation was quantified using a fluorescent probe, MitoSOX, imaged by confocal microscopy with a 60x objective. Scale bars depict 50 microns. [E] Quantification of mROS relative to Ctl cells in Ctl or KD NRAMs. n=62 cells. Statistical analyses were performed using Student’s t-test [B, C, E] to study differences between groups. Data are expressed as mean ± SEM.

Direct PITX2 transcriptional targets include electron transport chain complex I and IV which may generate excessive reactive oxygen species under mitochondrial dysfunction(Moris et al., 2017), and oxidative stress is a known pathological mechanism of atrial fibrillation(Jeong et al., 2012; Sovari, 2016; Xie et al., 2015). Therefore, we assessed mitochondrial-specific ROS (mitoROS) production using MitoSOX, a fluorescent mitoROS probe (Figure 2E). We observed elevated mitoROS in *Pitx2c*-deficient cardiomyocytes. All together, these data indicate that PITX2C transcriptionally regulates oxidative metabolism, and the loss of this regulatory network impairs mitochondrial dysfunction and subsequently leads to ROS accumulation.

### 3.3 *Pitx2c*-deficiency alters intracellular calcium cycling and arrhythmic oscillations

Calcium ions play a crucial role in cardiac excitation-contraction coupling with intracellular calcium oscillations during systole and diastole. PITX2 has been shown to directly transcriptionally regulate calcium cycling genes(Y. Tao et al., 2014) (Table S3) and *Pitx2* insufficiency results in abnormalities in calcium release(Lozano-Velasco et al., 2016; Pérez-Hernández et al., 2016; Tarifa et al., 2023; Vicente et al., 2024) thereby increasing afterdepolarizations, a key arrhythmogenic mechanism. Spontaneous intracellular calcium cycling was assessed to characterize calcium kinetics and the presence of arrhythmia in *Pitx2c*-deficient NRAMs (Figure 3A). *Pitx2c*-deficient cardiomyocytes beat faster (Figure 3B) in comparison to control cardiomyocytes yet, are slower to flux calcium during systole (Figure 3C) and are slower to reuptake calcium during diastole (Figure 3D). Afterdepolarization events were classified as either early (EADs) (Figure 3E) and delayed (DADs) (Figure 3F) afterdepolarizations as inferred from calcium traces. We observe a higher frequency of both EADs (Figure 3H) and DADs (Figure 3I). Together, these data indicate the presence of arrhythmia susceptibility in *Pitx2c*-deficient cardiomyocytes as a potential consequence of calcium dyshomeostasis.

**Figure 3.**
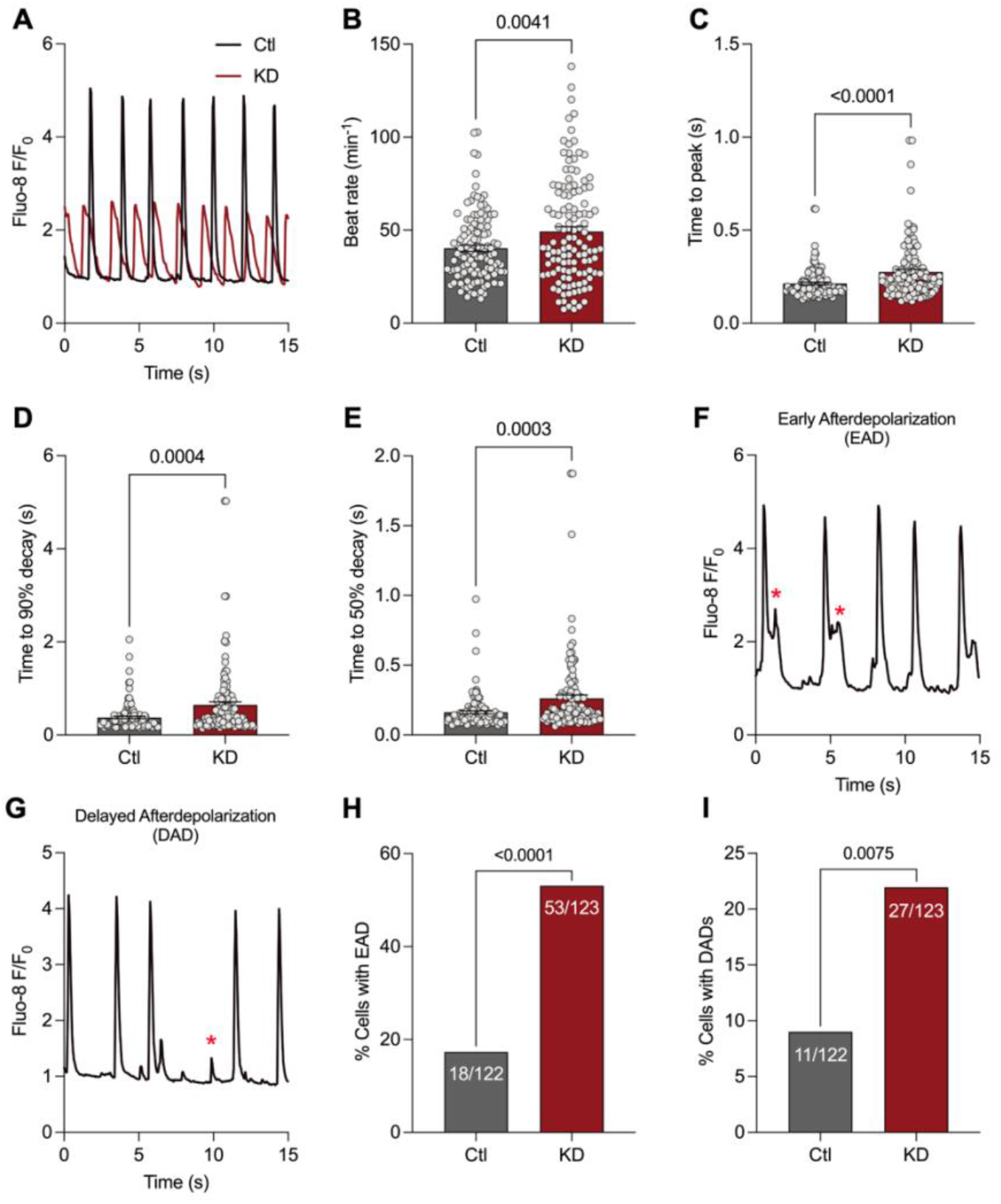
*Pitx2c* deficiency promotes ectopic beats through dysregulated intracellular calcium cycling. Spontaneous intracellular calcium dynamics were assessed in neonatal rat atrial myocytes (NRAMs) transfected with non-targeting (Ctl) or *Pitx2c* (KD) siRNA using a fluorescent calcium indicator, Fluo-8. [A] Representative time course of Fluo-8/baseline fluorescence (Fluo-8/F_0_) reporting rises in cytosolic calcium. Rate kinetics were quantified including [B] beat rate, [C] time to peak, [D] time to 90% decay, and [E] time to 50% decay. Early [F] and delayed [G] afterdepolarizations were classified via calcium traces and quantified [H,I]. Statistical analyses were performed using Student’s t-test [B-F] or Fisher’s exact test [G, H] to study differences between groups. Ctl: n=122 cells, KD: n=123. Data are expressed as means ± SEM.

### 3.4 *Pitx2c*-deficiency alters sarcomere structure and Titin protein localization

Structural remodelling occurs during AF pathogenesis; however, an underlying cardiomyopathy has also been suggested to increase cardiac arrhythmia susceptibility. *pitx2c-*deficient zebrafish display disorganized sarcomeres with absence of M-band structures(Collins et al., 2019). We initially characterized cellular morphology and observed an increase in cell size (Figure S2A), and indications of reduced cell elongation (Figure S2B, S2C). To further characterize AF-associated phenotypes present in *Pitx2c*-deficient NRAMs, we assessed subcellular sarcomere assembly using an antibody against Myomesin, an M-band protein (Figure 4A). *Pitx2c*-deficient cardiomyocytes had a shorter sarcomere length (Figure 4B) with no changes in sarcomere organization (Figure 4C). In contrast to the zebrafish model, we observe an increase in the density of sarcomere M-bands (Figure 4D). These data together may indicate a decrease in atrial contractility in relation to the decrease the sarcomere length is compensated by an increase in the overall density of sarcomeres.

**Figure 4.**
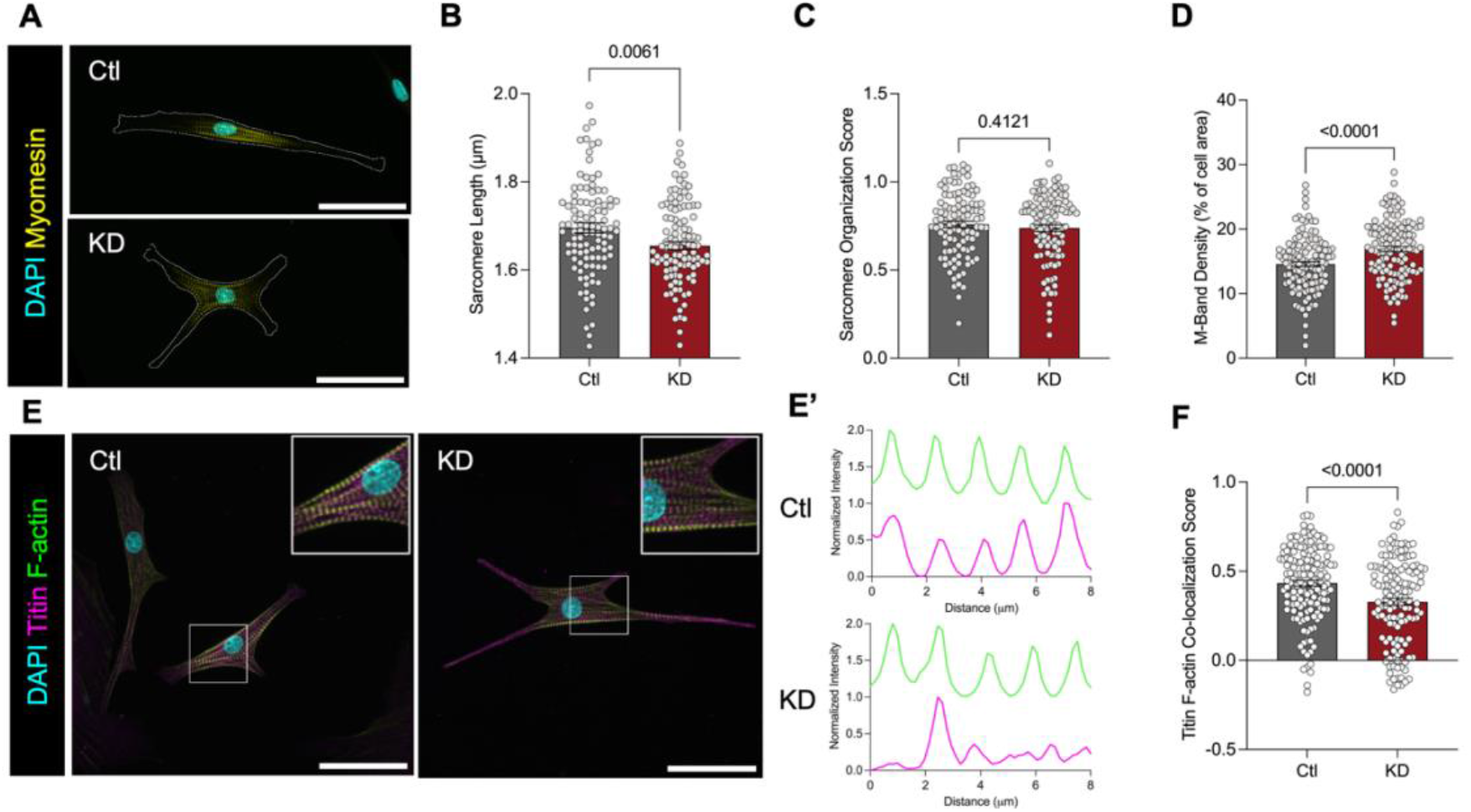
Sarcomeric proteins, Myomesin and Titin, are regulated by PITX2C. Sarcomere structure was assessed in neonatal rat atrial myocytes (NRAMs) transfected with non-targeting (Ctl) or *Pitx2c* (KD) siRNA. [A] Confocal microscopy of immunofluorescently labelled Myomesin (M-band), F-actin (Z-disk), and DAPI (nuclei). Scale bars depict 50 microns. Quantification of [B] sarcomere length [C] sarcomere organization and [D] M-band density (as a % of total cell area) [A,B] Ctl: n=105 cells, KD: n=106 cells. [D] Ctl: n=124 cells, KD: n=126 cells [E] Confocal microscopy of immunofluorescently labelled Titin, F-actin, and DAPI. Scale bars depict 50 microns and [E’] representation of co-localization of Titin and F-actin signal along one myofibril. [F] Quantification of Titin and F-actin co-localization. Ctl: n=154 cells, KD: 140 cells. Statistical analyses were performed using Student’s t-test [B-D, F] to study differences between groups. Data are expressed as mean ± SEM.

It is of note to mention that transcriptomic analysis reveal myogenesis as one of the top dysregulated pathways in *Pitx2c*-deficiency (Figure 1C) and a number direct PITX2 transcriptional targets exist at the thick/thin filaments (*Myh6, Myh7b, Myh8, Tnni1, Tnni3k, Tnnt2, Tpm2, Tpm4*), Z-disk (*Neb, Pdlim5, Pdlim4, Capzb, Fmn2, Lmod1, Tmod1*), and M-band (*Ampd2, Obscn, Capn3*) (Table S3). We additionally performed immunocytochemistry for Titin, a crucial protein within the sarcomere in maintaining cardiomyocyte elasticity (Figure 4E). We observed that *Pitx2c*-deficient cardiomyocytes display Titin mis-localization within the sarcomere (Figure 4F). We quantified total Titin levels via qPCR targeting exons 49-50 and identified an upregulation of total Titin mRNA expression (Figure S2D), suggesting that altered Titin levels may contribute its mislocalization. We also considered the splicing factor RNA-binding motif protein 20 (Rbm20), which acts to alternatively splice Titin. Mutations in *Rbm20* have been associated with arrhythmogenic cardiomyopathies with erroneous splicing resulting in high expression of the fetal N2BA Titin isoform(Van Den Hoogenhof et al., 2018; Y. Zhang et al., 2023). Notably, ChIP-seq analysis identifies *Rbm20* as a direct transcriptional target of PITX2 (Table S3).

### 3.5 N-acetylcysteine reduces oxidative stress and arrhythmia-associated phenotypes in *Pitx2c*-deficiency

There are several ways by which ROS may induce cardiac arrhythmia, including by oxidizing calcium cycling or sarcomere proteins, which are in line with some of the phenotypes we observe in *Pitx2c*-deficient atrial cardiomyocytes. To evaluate ROS as a mechanism by which the loss of *Pitx2c* drives arrhythmogenesis, we treated control and *Pitx2c*-deficent cells with 1mM NAC, a potent ROS scavenger, for 24 hours. NAC reduced mitochondrial-derived ROS accumulation (Figure 5A) in *Pitx2c*-deficient atrial cardiomyocytes. In addition, reducing ROS restored PITX2C-dependent alterations in calcium cycling parameters including beat rate (Figure 5B) and time to decay (Figure 5D, 5E). However, while NAC treatment reduces the time to peak (Figure 5C), it was unable to restore the prolongation in calcium release (Figure 5C) back to control levels. Despite this, the frequency of both early and delayed afterdepolarizations was reduced (Figure 5G, 5H). We additionally assessed Titin localization following NAC treatment and observe Titin is relocalized to the Z-disk. Together, these data suggest that PITX2C loss results in ROS accumulation that promotes abnormal calcium cycling and sarcomere instability, strengthening the hypothesis that PITX2C is a critical regulator of mitochondrial function and oxidative stress in atrial cardiomyocytes.

**Figure 5.**
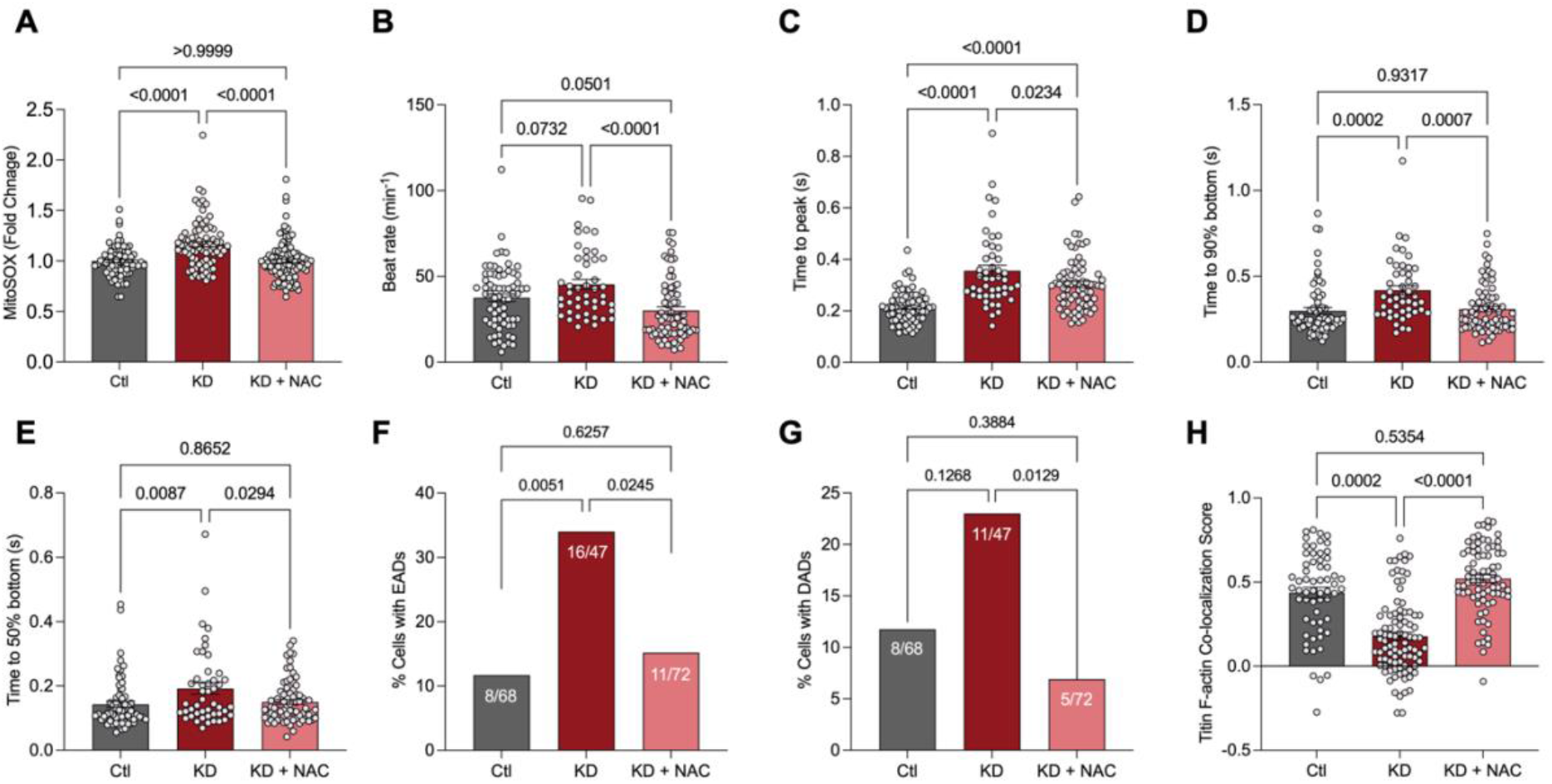
Antioxidant treatment reduces deleterious phenotypes in *Pitx2c* deficient atrial cardiomyocytes. Neonatal rat atrial myocytes (NRAMs) transfected with non-targeting (Ctl) or *Pitx2c* (KD) siRNA were treated with 1mM N-acetylcysteine (NAC) for 24 hours. [A] Quantification of mitochondrial specific reactive oxygen species using MitoSOX relative to Ctl cells in Ctl or KD NRAMs. Ctl: 76 cells, KD: 79 cells, KD + NAC: 96 cells. Intracellular calcium rate kinetics were quantified including [B] beat rate, [C] time to peak, [D] time to 90% decay, and [E] time to 50% decay. Early [F] and delayed [G] afterdepolarizations via calcium traces and quantified. Ctl: n=68 cells, KD: n=47, KD + NAC: 72 cells. [H] Quantification of Titin and F-actin co-localization. Ctl: 60 cells, KD: 91 cells, KD + NAC: 76 cells. Statistical analyses were performed using Student’s t-test [A-E, H] or Fisher’s exact test [F,G] to study differences between groups. Data are expressed as mean ± SEM.

## 4 DISCUSSIONS

### 4.1 Main findings

In this study, our data reveal reactive oxygen species as an early mechanism of intracellular calcium handling and sarcomere structure downstream of PITX2. We identify altered oxidative metabolism and elevations in reactive oxygen species accumulation in *Pitx2c* deficient neonatal rat atrial myocytes. Treatment with NAC, reduces AF-associated phenotypes including afterdepolarizations and Titin localization to the sarcomere. In addition, we report a novel interaction between PITX2 and Titin, an integral sarcomere protein.

### 4.2 PITX2 regulates atrial metabolism

We report an evolving role of PITX2 in regulating atrial metabolism. In this study, we observe *Pitx2c*-deficient cardiomyocytes display decreased basal and maximal oxidative phosphorylation as well as decreased ATP production. These findings are consistent with *pitx2c* deficient zebrafish, murine models of *Pitx2* insufficiency(L’honoré et al., 2014; Li et al., 2018; G. Tao et al., 2016), *PITX2* deficient human induced pluripotent stem cells (hiPSC)(Benzoni et al., 2023; Reyat et al., 2024), and AF patient derived left atrial tissue(Reyat et al., 2024). In addition, it is becoming increasingly recognized that decreased oxidative phosphorylation is a key finding in atrial fibrillation(Emelyanova et al., 2016; Kanaan et al., 2019) which may contribute energy deficits, oxidative stress, and structural and electrical remodelling that causes arrhythmia. We identify amongst the top differentially expressed genes, direct PITX2 transcriptional targets including electron transport chain complex I and IV subunits which may explain the accumulation of ROS(Moris et al., 2017). A limitation to note is that the ChIP-seq experiment was previously performed in 12-week-old mouse heart tissue which are both different species and at a different developmental stage than the neonatal rat model used within this study.

Metabolic deficiency is a key finding in arrhythmia and heart failure with a decrease in oxidative metabolism and a shift towards glycolysis as an initial cardioprotective mechanism(Lopaschuk et al., 2021). This metabolic switch has been demonstrated in *PITX2* deficient atrial hiPSCs(Reyat et al., 2024). Although we observe transcriptional changes in glycolysis, we observe no changes in basal glycolytic function. Potentially, as our study leverages neonatal rat cardiomyocytes that are already undergoing a dynamic developmental metabolic switch from glycolysis to fatty acid metabolism(Piquereau & Ventura-Clapier, 2018), the loss of PITX2 in NRAMs, rather than decreasing glycolytic function, hinders this metabolic switch. Therefore, without PITX2 that regulates atrial metabolism, neonatal rat cardiomyocytes remain utilizing glycolysis rather than fatty acid metabolism, an abundant energy producing source. This presents an area for further exploration to understand how PITX2 regulates post-natal glucose and fatty acid import and utilization in atrial cardiomyocyte development.

### 4.3 PITX2 regulates calcium cycling and sarcomere structure

The role of PITX2 in transcriptionally regulating calcium handling genes and in turn cardiac rhythm is well established(Lozano-Velasco et al., 2016; Tarifa et al., 2023; Vicente et al., 2024). Surprisingly, we fail to observe significant dysregulation of genes encoding cardiac calcium handling genes including L-type calcium channels (*Cacna1d*), cardiac ryanodine receptors (*Ryr2*), and SERCA2a (*Atp2a2*). However, we observe functional deficits in calcium handling, such as alterations in calcium transients, including both the time to peak and decay which are indicative of RyR2 and SERCA2a calcium flux, respectively. Therefore, we hypothesize that oxidation of these proteins drive the alterations in calcium flux as treatment with N-acetylcysteine restores calcium transient parameters and afterdepolarizations(Balderas-Villalobos et al., 2013; Boraso & Williams, 1994; Marengo et al., 1998; Plummer et al., 2015; Salama et al., 2000; Sun et al., 2008; Xu et al., 1998). However, as we observe that the time to peak remains elevated, there may be an irreversible post-translational modification of the RyR2 leading to its sustained activation(Xu et al., 1998). Further, it has been previously shown that augmented RyR2 activity modulates mitochondrial calcium handling and promotes mitoROS emission resulting in a pro-arrhythmic feedback cycle(Hamilton et al., 2020). A limitation of this study is that we do not assess the redox state or modification of the receptor (S-nitrosylation, S-glutathionylation, disulfide oxidation) directly. Nonetheless, oxidative stress appears to promote arrhythmic mechanisms and reducing oxidative stress reduces arrhythmia.

Titin is an integral sarcomere protein, known for its role in maintaining cardiomyocyte elasticity. Truncating mutations in Titin have been largely implicated in dilated cardiomyopathies(Herman et al., 2012) which subsequently increases the risk of developing arrhythmia. Jiang and Ly et al. demonstrated that a 9-amino acid deletion from Titin’s A-band in zebrafish and hiPSC atrial-cardiomyocytes results in electrophysiological remodelling that increases AF susceptibility(Jiang et al., 2024). This study suggests oxidation of Titin downstream of PITX2 is a cardiomyopathic component that coincides with AF susceptibility. Oxidation of Titin protein is known to increase or decrease compliance depending on site of oxidation which may impair diastolic function(Loescher et al., 2020). However, how PITX2 regulates Titin protein localization to the sarcomere remains unanswered. One potential mechanism is through high levels of ROS and oxidation of the Titin protein activates protein quality control mechanisms(Rudolph et al., 2019) that result in the degradation of the protein, a mechanism specific to Titin still largely unknown. Or, supported by our transcriptomic data PITX2 regulates neonatal myogenesis and may transcriptionally regulate Titin isoform switching from a passive N2BA neonatal isoform to a stiff adult N2B isoform(Warren et al., 2004). This may be further supported by *Rbm20*, a Titin splicing factor, being identified as a direct transcriptional target of PITX2. In addition, PITX2 may regulate other key sarcomere proteins which alter Titin anchoring and therefore results in mis-localization. Although incompletely understood, this interaction presents an exciting avenue for further studies.

It is important to consider the relationship between Titin protein function and intracellular calcium dynamics. In particular, the PEVK segment, rich in E (glutamic acid) residues, binds calcium with high affinity, increasing passive tension with increasing levels of calcium. We identify changes in intracellular calcium dynamics, highlighting a prolonged calcium cycle, which may in part alter the passive forces produced by Titin and other sarcomere components. In addition, direct transcriptional targets of PITX2 lie in Ca^2+^/calmodulin-dependent protein kinase and S100 calcium binding protein which have been demonstrated to interact with Titin at the N2B and PEVK regions(Hamdani et al., 2013; Hidalgo et al., 2013; Perkin et al., 2015) and N2A region(Apel et al., 2025; Yamasaki et al., 2001), respectively.

### 4.4 Reactive oxygen species in atrial fibrillation

Current anti-arrhythmic drugs commonly used in the treatment of AF act by blocking B-adrenergic receptors or ion channels(Joglar et al., 2024). While these management strategies are beneficial, they are found to have limited long-term efficacy and induce pro-arrhythmic effects(Heijman et al., 2018). Therefore, due to the clear implication of oxidative stress in AF pathogenesis, understanding the molecular mechanisms of AF and identifying novel therapeutic targets is necessary. In this study, we identify treatment with NAC restores key components of cardiomyocyte function by reducing levels of ROS where a *PITX2* genetic predisposition is present. However, clinical implementation of NAC as a treatment for AF has been controversial. Results vary amongst clinical trials in administering NAC as a preventative measure of postoperative AF while some studies suggest reduced incidence, others suggest no benefit(Violi et al., 2014). These differences in outcomes may be due in part to the incomplete understanding of ROS and the vast number of intracellular targets and consequences. In addition, how a genetic component contributes to the drug interactions and efficacy is incomplete. However, we shed insight into the efficacy of NAC in PITX2C deficiency, the most significant genetic risk associated with AF. The long-term efficacy of antioxidants in the treatment of other AF subtypes remains incomplete.

### 4.4 The potential role of PITX2 in the link between inflammation and atrial fibrillation

Although outside of the scope of this study, inflammation is a key contributor to reactive oxygen species in AF pathogenesis. We identify dysregulated pathways in inflammation, TNF-α signaling via NF-KB, and the complement system suggesting that PITX2 transcriptionally activates inflammatory signaling (Figure 1C). Concurrent with these findings, it has been previously identified that inflammatory gene expression was upregulated in a mouse model with a deletion of an intronic CCCTC-binding factor (CTCF) site in the *Pitx2* gene leading to reduced *Pitx2* expression(M. Zhang et al., 2019). The precise downstream mechanisms of PITX2 in regulating inflammation in AF remains unclear. However, ROS and inflammation may act cooperatively to exacerbate atrial electrical and structural remodeling that facilitates AF(Mittal et al., 2014; Nso et al., 2021). Zhou and Liu et al. previously reported mouse models of diabetes mellitus with increased AF inducibility that was primarily driven macrophage IL-1B through mitoROS modulation of the RyR2(Zhou et al., 2024). The mechanisms of ROS and inflammation in *Pitx2c* deficiency warrants further investigation.

## ABBREVIATIONS

AF: Atrial fibrillation
PITX2: paired-like homeodomain transcription factor 2
NRAM: neonatal rat atrial myocyte
ROS: reactive oxygen species
mitoROS: mitochondrial reactive oxygen species
EAD: early afterdepolarization
DAD: delayed afterdepolarization
NAC: N-acetylcysteine
OCR: oxygen consumption rate
FCCP: Carbonyl cyanide-4 (trifluoromethoxy) phenylhydrazone
PER: proton efflux rate

## 7 AUTHOR CONTRIBUTIONS

*Conceptualization*: AK, MMC

*Conducted experiments:* AK, SG

*Data analysis:* AK, SG, FSV, MMC

*Writing-original draft:* AK, MMC

*Writing-review and editing:* AK, SG, FSV, MMC

Supervision: MMC

## 8 ACKNOWLEDGEMENTS

The authors are grateful to undergraduate students Jared Stevenson and Kenzie Byers, and all members of the Collins Lab, both past and present. The authors are grateful to Betty Chow-Lockerbie for technical assistance with Seahorse experiments.

## 9 DATA AVAILABILITY

The RNA-seq dataset presented in this study have been deposited to The Gene Expression Omnibus (GEO) under accession number GSE325239.

