## Supplemental Figures for "PITX2C Deficiency Promotes Arrhythmogenic Remodeling via Oxidative Stress in Atrial Myocytes"

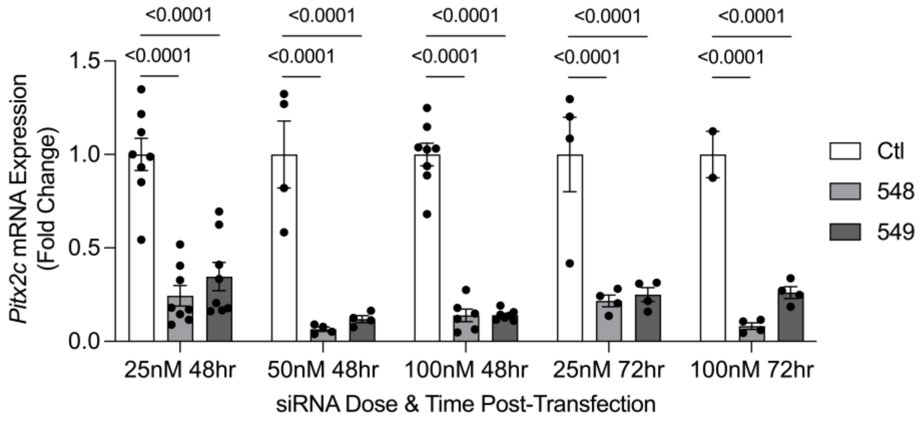


**Figure S1.** Neonatal rat atrial myocytes transfected with non-targeting (Ctl) siRNA or one of two *Pitx2c* (548, 549) siRNAs at various doses. RT-qPCR was performed at 48- and 72-hours post-transfection and *Pitx2c* mRNA levels were normalized to B-actin. Statistical analyses were performed using 2-way ANOVA followed by Tukey's multiple comparison test. Data are expressed as mean ± SEM.

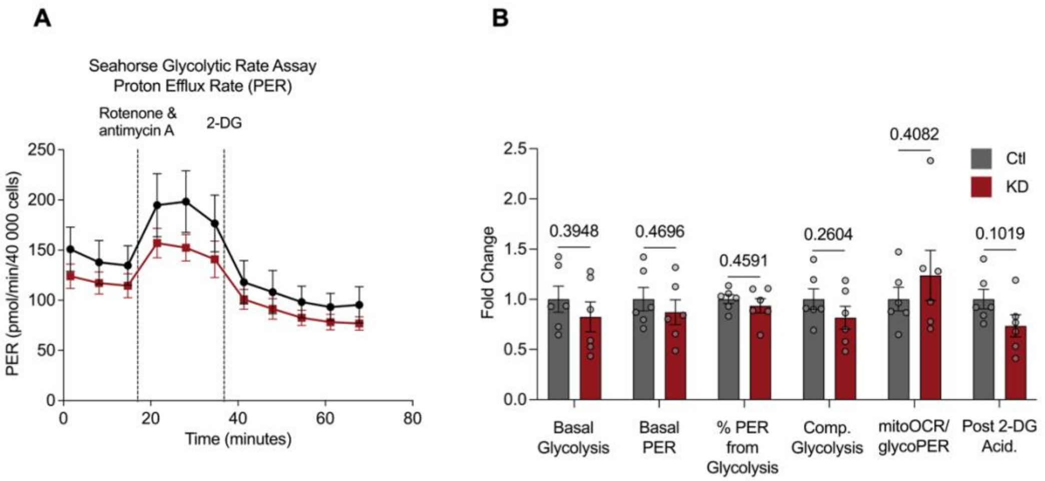


**Figure S2. *Pitx2c* deficiency does not impair glycolytic function.** Glycolysis was assessed in neonatal rat atrial myocytes (NRAMs) transfected with non-targeting (Ctl) or *Pitx2c* (KD) siRNA using the Seahorse XFp Glycolytic Rate assay. [A] Representative proton efflux rate (PER) trace is shown with injections of rotenone & antimycin A and 2-deoxyglucose (2-DG). [B] Quantification of glycolytic function relative to Ctl cells including basal glycolysis, basal PER, % PER from glycolysis, compensatory glycolysis, mitochondrial oxygen consumption rate (mitoOCR)/glycolysis PER (glycoPER), and post 2-DG acidification (post 2-DG acid.). n=6 from 2 independent experiments. Statistical analyses were performed using Student's t-test [B, C, E] to study differences between groups. Data are expressed as mean ± SEM.

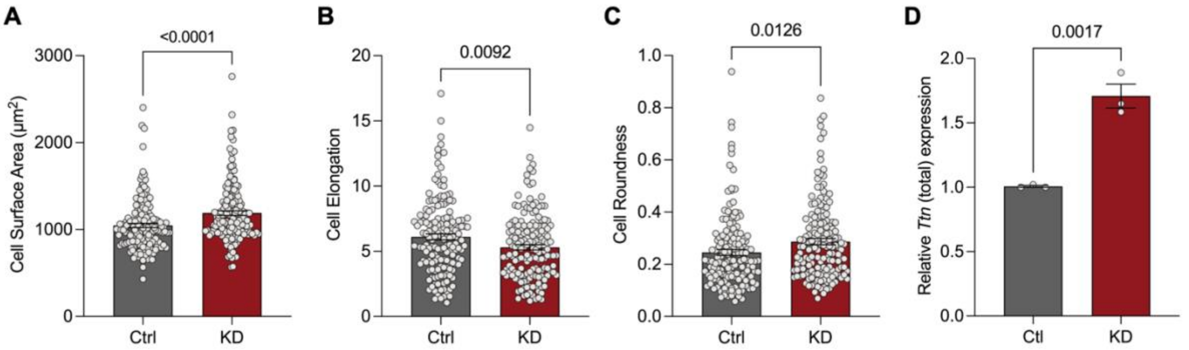


**Figure S3. *Pitx2c* deficiency results in an increase in cell surface area and overall morphological alterations.** Cell morphology was assessed in fluorescently labelled neonatal rat atrial myocytes (NRAMs) transfected with non-targeting (Ctl) or *Pitx2c* (KD) siRNA stained for F-actin. [A] Cell surface area, [B] cell elongation, and [C] cell roundness were quantified. n=310-340 cells from 3 independent experiments. Data are expressed as means ± SEM. Statistical analyses performed using Student's t-test to study differences between groups. RT-qPCR of quantify [D] total Titin (exon 49-50) mRNA levels. Total Titin levels were normalized to B-actin. n=3. Data are expressed as means ± SEM. Statistical analyses performed using Student's t-test to study differences between groups.

**Table S1.** Primers used for RT-qPCR.

| Gene | Forward oligo | Reverse oligo |
|---|---|---|
| *Pitx2c* | CGATACGTCCAGCCCTGAAG | GTTGCCGCTTCTTCTTGGAC |
| *Actb* | AGGCCAACCGTGAAAAGATG | ACCAGAGGCATACAGGGACAA |
| *mt-Co1*[58] | TGGAGGCTTCGGAAACTGACT | ATGGGCTAGGTTTCCGGCTAA |
| *Polg* | ATTGGTGATCTGCAGTGCCAG | CAGTCGTGTGAATTGGGTGGT |
| *Ttn* (total)[59] | CCAAGCTCACTGTGGGAGAAA | GCTACTTCCAAGGGCTCAATTC |

**Table S2.** Electron transport chain (ETC)-related genes identified from ChIP-seq[5].

| **Gene** | ***ETC component*** |
|---|---|
| *Ndufaf6* | Complex I |
| *Ndufaf8* | Complex I |
| *Ndufb1* | Complex I |
| *Ndufb11b* | Complex I |
| *Ndufb3* | Complex I |
| *Ndufb9* | Complex I |
| *Ndufs6b* | Complex I |
| *Ndufs7* | Complex I |
| *Ndufv1* | Complex I |
| *Ndufv3* | Complex I |
| *Uqcc1* | Complex III |
| *Uqcc2* | Complex III |
| *Uqcc4* | Complex III |
| *Uqcc5* | Complex III |
| *Coa3* | Complex IV |
| *Coa8* | Complex IV |
| *Cox7b2* | Complex IV |
| *Ndufa4* | Complex IV |
| *Atp5mj* | ATP synthase |

**Table S3.** Select calcium and sarcomere genes identified from ChIP-seq[5].

| ***Gene*** | ***Description*** |
|---|---|
| *Cacna1e* | Calcium channel, voltage-dependent, R type, alpha 1E subunit |
| *Camk1d* | Calcium/calmodulin-dependent protein kinase IDI |
| *Cacna1b* | Calcium channel, voltage-dependent, N type, alpha 1B subunit |
| *Cacnb4* | Calcium channel, voltage-dependent, beta 4 subunitI |
| *S100a3* | S100 calcium binding protein A3 |
| *S100a10* | S100 calcium binding protein A10 (calpactin) |
| *Camk2d* | Calcium/calmodulin-dependent protein kinase II, delta |
| *Efcab14* | EF-hand calcium binding domain 14 |
| *Cabs1* | Calcium binding protein, spermatid specific 1 |
| *Ppef2* | Protein phosphatase, EF hand calcium-binding domain 2 |
| *Cacna1c* | Calcium channel, voltage-dependent, L type, alpha 1C subunit |
| *Cracr2a* | Calcium release activated channel regulator 2A |
| *Cacna2d2* | Calcium channel, voltage-dependent, alpha 2/delta subunit 2 |
| *Calhm6* | Calcium homeostasis modulator family member 6 |
| *Mcu* | Mitochondrial calcium uniporter |
| *Calcoco2* | Calcium binding and coiled-coil domain 2 |
| *Efcab3* | EF-hand calcium binding domain 3 |
| *Smoc1* | SPARC related modular calcium binding 1 |
| *Slc24a4* | Solute carrier family 24 (sodium/potassium/calcium exchanger), member 4 |
| *Cacng2* | Calcium channel, voltage-dependent, gamma subunit 2 |
| *Efcab6* | EF-hand calcium binding domain 6 |
| *Cacnb3* | Calcium channel, voltage-dependent, beta 3 subunit |
| *Myh7b* | Myosin, heavy chain 7B, cardiac muscle, beta |
| *Myh8* | Myosin, heavy polypeptide 8, skeletal muscle, perinatal |
| *Myh6* | Myosin, heavy polypeptide 6, cardiac muscle, alpha |
| *Tnni1* | Troponin I, skeletal, slow 1 |
| *Tnni3k* | TNNI3 interacting kinase |
| *Tnnt2* | Troponin T2, cardiac |
| *Tpm2* | Tropomyosin 2, beta |
| *Tpm4* | Tropomyosin 4 |
| *Neb* | Nebulin |
| *Pdlim5* | PDZ and LIM domain 5 |
| *Pdlim4* | PDZ and LIM domain 4 |
| *Capzb* | Capping actin protein of muscle Z-line subunit beta |
| *Fmn2* | Formin 2 |
| *Lmod1* | Leiomodin 1 (smooth muscle) |
| *Tmod1* | Tropomodulin 1 |
| *Ampd2* | Adenosine monophosphate deaminase 2 |
| *Obscn* | Obscurin, cytoskeletal calmodulin and titin-interacting RhoGEF |
| *Capn3* | Calpain 3 |
| *Rbm20* | RNA binding motif protein 20 |
